# Discovery and characterization of non-canonical E2 conjugating enzymes

**DOI:** 10.1101/2023.03.05.531151

**Authors:** Syed Arif Abdul Rehman, Elena Di Nisio, Chiara Cazzaniga, Odetta Antico, Axel Knebel, Clare Johnson, Frederic Lamoliatte, Rodolfo Negri, Miratul Muqit MK, Virginia De Cesare

## Abstract

E2 conjugating enzymes (E2s) play a central role in the enzymatic cascade that leads to the attachment of ubiquitin to a substrate. This process, termed ubiquitylation is fundamental for maintaining cellular homeostasis and impacts almost all cellular process. By interacting with multiple E3 ligases, E2s direct the ubiquitylation landscape within the cell. Since its discovery, ubiquitylation has been regarded as a post-translational modification that specifically targets lysine side chains (canonical ubiquitylation). We used MALDI-TOF Mass Spectrometry to discover and characterize a family of E2s that are instead able to conjugate ubiquitin to serine and/or threonine. We employed protein modelling and prediction tools to identify the catalytic determinants that these E2s use to interact with ubiquitin as well as their substrates. Our results join a stream of recent literature that challenges the definition of ubiquitylation as an exquisitely lysine-specific modification and provide crucial insights into the missing E2 element responsible for non-canonical ubiquitylation.

**Teaser:** E2 conjugating enzymes (E2s) play a fundamental role in the attachment of ubiquitin to its substrate. Most E2s can form an isopeptide bond between the ubiquitin C- terminus and a lysine present on the substrate. We identified a family of E2s, UBE2Q1 and UBE2Q2, able to target amino acids other than lysine. Currently nothing is known about their mechanism of action and what substrates they are targeting, even though genetic ablation of UBE2Q1 produce substantial infertility in mice. Here we answer the question about what the key residues beneath their peculiar activity are. We discovered that UBE2Q1 target the lysine-free cytoplasmic domain of the Golgi resident protein Beta-1,4-galactosyltransferase 1, providing an interesting precedent for the role of non-canonical ubiquitylation in eukaryotic cells.

## Introduction

Attachment of one or more ubiquitin molecules to a substrate requires the sequential activity of an E1 activating enzyme, an E2 conjugating enzyme and an E3 ligase. This process, named ubiquitylation (or ubiquitination) plays a major role in various pathways during cell life and death, including but not limited to cell division and differentiation, response to environmental stress, immune response, DNA repair, and apoptosis ^1–6^. The human genome encodes around 40 E2s ^7^ and more than 700 E3 ligases ^8, 9^. E3 ligases are divided into subfamilies depending on the presence of either a RING (Really Interesting New Gene) or HECT (Homologous to the E6AP Carboxyl Terminus) domain ^10^. RING E3 ligases represent the vast majority of known E3s ^8^ and they represent essential activators that facilitate the direct transfer of ubiquitin from the E2s to the substrate by decreasing the Km and increases Kcat for both their substrates: Ub-loaded E2 and the protein to be modified. Besides the activating role of the RING E3 ligases. E2 conjugating enzymes possess the catalytic determinants that direct the transfer of ubiquitin to the substrate and govern both the type of ubiquitin linkage and the extent of ubiquitin modification ^11, 12^. E2s that functionally interact with RING E3 ligases have intrinsic reactivity toward lysine, the canonical ubiquitylation target. However, other non-canonical, hydroxyl-containing amino acid and biomolecules, such as serine, threonine, sugars, and the bacterial liposaccharide (LPS), have been found to be also targeted via E3-mediated ubiquitylation ^13–23^ and by the ubiquitin-like protein urm1^24^. The isopeptide bond formed between the ubiquitin C-terminus and the amine present in the lysine side chain is very stable over a range of temperature and pH. On the other hand, the ester bond formed between the ubiquitin C-terminus and the hydroxyl group present in non-canonical targets is hydrolysed in mild basic conditions and relative low temperatures. Because of the intrinsically labile nature of the ester bond and the lack of high-resolution dedicated analytical tools, the identification of ubiquitin conjugating enzymes able to target residues other than lysine remains challenging. Here we develop a MALDI-TOF Mass Spectrometry based assay to systematically interrogate ubiquitin conjugating enzymes for their ability to ubiquitylate lysine and other non-canonical residues. We identify a new family of E2s (UBE2Qs) that ubiquitylates non-canonical residues such as serine, threonine but also other biomolecules, including variously complex sugars. The UBE2Q family set themselves apart from canonical E2s in several aspects, they do not possess the canonical Histidine-Proline- Asparagine (HPN) catalytic triad that characterizes canonical E2s and have an extended N- terminus. We used Alpha Fold ^25^ and COOT ^26^ software to generate a structural model and predict the catalytic determinants which were validated by mutational and biochemical analyses. Because E2 acts upstream of E3 in the ubiquitylation cascade and can interact with multiple RING E3 ligases ^27^, E2s have a larger range of substrates compared to the more specific E3 ligases. We therefore anticipate that the discovery of new E2s with non-canonical activity will have profound and wide- ranging impacts on the ubiquitylation landscape and, consequently, on biological processes.

## Materials and Methods

### E1 Activating enzyme and E2s conjugating enzymes expression and purification

Human recombinant 6His-tagged UBE1 was expressed in and purified from Sf21 cells using standard protocols. Human E2s were all expressed as 6His-tagged fusion proteins in BL21 cells and purified via their tags using standard protocols as previously described ^28^. Briefly, BL21 DE3 codon plus cells were transformed with the appropriate constructs (see Table1), colonies were picked for overnight cultures, which were used to inoculate 6 x 1L LB medium supplemented with antibiotics. The cells were grown in Infors incubators, whirling at 200 rpm until the OD600 reached 0.5 – 0.6 and then cooled to 16°C – 20°C. Protein expression was induced with typically 250 μM IPTG and the cells were left over night at the latter temperature. The cells were collected by centrifugation at 4200 rpm for 25min at 4°C in a Beckman J6 centrifuge using a 6 x 1 L bucket rotor (4.2). The cells were resuspended in ice cold lysis buffer (50 mM Tris-HCl pH 7.5, 250 mM NaCl, 25 mM imidazole, 0.1 mM EGTA, 0.1 mM EDTA, 0.2 % Triton X-100, 10 μg/ml Leupeptin,

1 mM PefaBloc (Roche), 1mM DTT) and sonicated. Insoluble material was removed by centrifugation at 18500 xg for 25 min at 4°C. The supernatant was incubated for 1 h with Ni-NTA- agarose (Expedeon), then washed five times with 10 volumes of the lysis buffer and then twice in 50 mM HEPES pH 7.5, 150 mM NaCl, 0.015% Brij35, 1 mM DTT. Elution was achieved by incubation with the latter buffer containing 0.4M imidazole or by incubation with Tobacco Etch Virus (TEV) protease (purified in house). The proteins were buffer exchanged into 50 mM HEPES pH 7.5, 150 mM NaCl, 10% glycerol and 1 mM DTT and stored at -80°C.

### Screening of E2 conjugating enzymes activity by MALDI-TOF MS

23 recombinant E2 conjugating enzymes (see Table 1) were expressed and adjusted to a final concentration of 2 µM final into a mixture containing UBE1 (200 nM.), 2 mM ATP, 20 mM MgCl2, 2 mM TCEP and 1X phosphate buffer (PBS, pH 7.5). 5 µL of enzymatic mixture was then dispensed using an electronic 16 multichannel pipet into a Lowbind 384 Eppendorf plate. Stock solution of Acetyl-Lysine (Ac-K), Acetyl-Serine (Ac-S), Ac-Threonine (Ac-T), glycerol and glucose were prepared at the final concentration of 500 mM and pH adjust to ∼7.5. 2 µL of Ac-K, c-S, Ac-T, glycerol or glucose were independently added to the enzymatic mixture. The reaction was started by adding 5 µL of ubiquitin (2 µM) in 1X PBS. The assay plates were covered with a self-adhesive aluminium foil and incubated at 30°C for the indicated time point(s) in an Eppendorf ThermoMixer C (Eppendorf) equipped with a ThermoTop 288 and a SmartBlock™ PCR 384. The reactions were stopped by adding 6% TFA supplemented with ^15^N Ubiquitin (2 µM). Samples were spotted on 1536 AnchorChip MALDI plate using Mosquito nanoliter pipetting system (TTP Labtech) as previously reported ^28–31^. Detection by MALDI-TOF/MS was also performed similarly to previously described ^29^

**Table 1.**
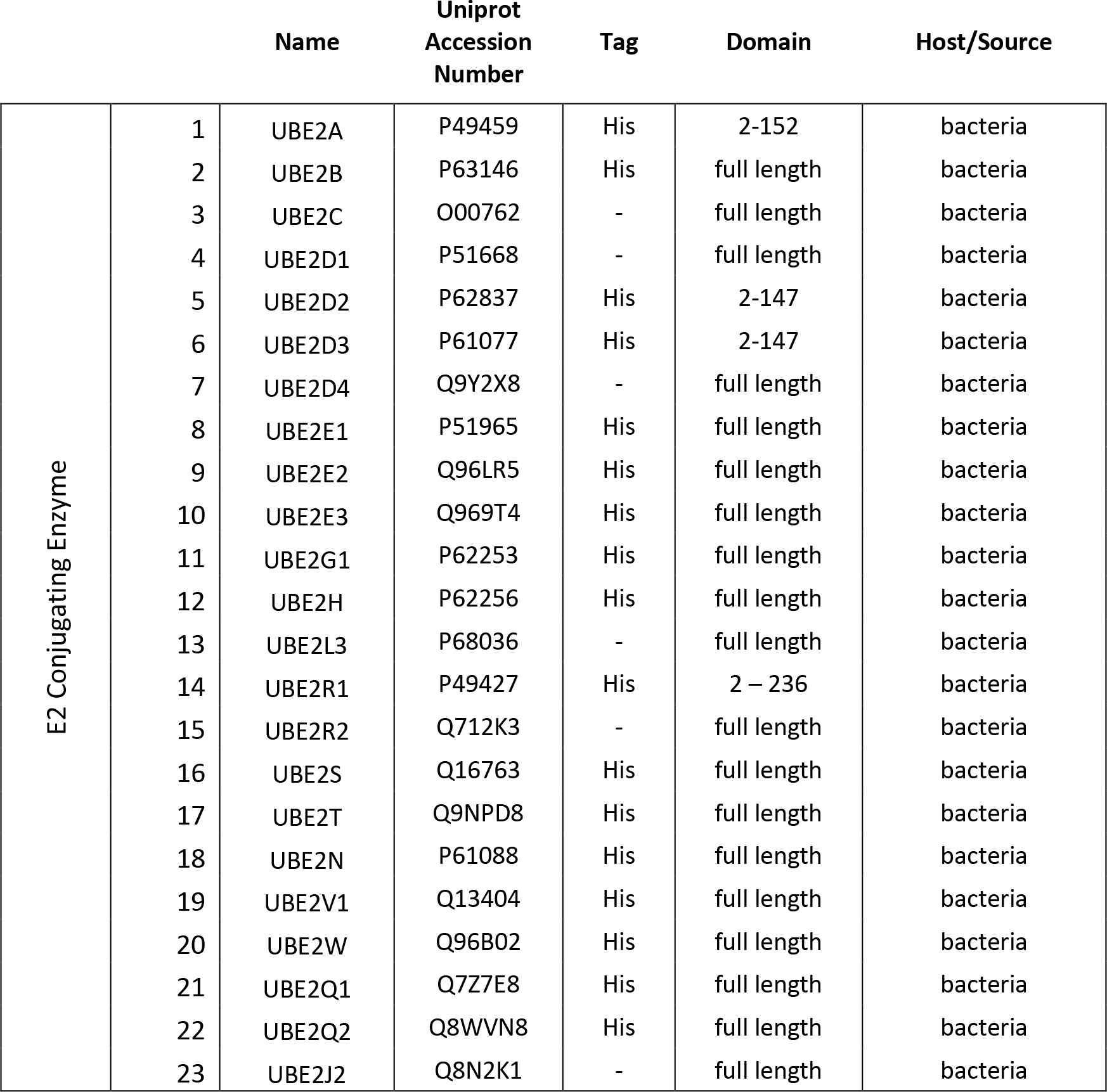
E2 conjugating enzymes in use in this study.

|  |  | Name | Uniprot<br>Accession<br>Number | Tag | Domain | Host/Source |
| --- | --- | --- | --- | --- | --- | --- |
| E2 Conjugating Enzyme | 1 | UBE2A | P49459 | His | 2-152 | bacteria |
|  | 2 | UBE2B | P63146 | His | full length | bacteria |
|  | 3 | UBE2C | O00762 | - | full length | bacteria |
|  | 4 | UBE2D1 | P51668 | - | full length | bacteria |
|  | 5 | UBE2D2 | P62837 | His | 2-147 | bacteria |
|  | 6 | UBE2D3 | P61077 | His | 2-147 | bacteria |
|  | 7 | UBE2D4 | Q9Y2X8 | - | full length | bacteria |
|  | 8 | UBE2E1 | P51965 | His | full length | bacteria |
|  | 9 | UBE2E2 | Q96LR5 | His | full length | bacteria |
|  | 10 | UBE2E3 | Q969T4 | His | full length | bacteria |
|  | 11 | UBE2G1 | P62253 | His | full length | bacteria |
|  | 12 | UBE2H | P62256 | His | full length | bacteria |
|  | 13 | UBE2L3 | P68036 | - | full length | bacteria |
|  | 14 | UBE2R1 | P49427 | His | 2 – 236 | bacteria |
|  | 15 | UBE2R2 | Q712K3 | - | full length | bacteria |
|  | 16 | UBE2S | Q16763 | His | full length | bacteria |
|  | 17 | UBE2T | Q9NPD8 | His | full length | bacteria |
|  | 18 | UBE2N | P61088 | His | full length | bacteria |
|  | 19 | UBE2V1 | Q13404 | His | full length | bacteria |
|  | 20 | UBE2W | Q96B02 | His | full length | bacteria |
|  | 21 | UBE2Q1 | Q7Z7E8 | His | full length | bacteria |
|  | 22 | UBE2Q2 | Q8WVN8 | His | full length | bacteria |
|  | 23 | UBE2J2 | Q8N2K1 | - | full length | bacteria |

. Briefly, all samples were acquired on a Rapiflex MALDI TOF mass spectrometer (Bruker Daltonics, Bremen, Germany) high resolution MALDI-TOF MS instrument with Compass for flexSeries 2.0 equipped with FlexControl Version 4.0, Build 48 and FlexAnalysis version 4.0, Build 14. Sample spectra were collected in automatic mode using AutoXecute, Bruker Daltonics, Fuzzy Control parameters were switched off, initial Laser Power set on “from Laser Attenuator” and accumulation parameters set to 4000 satisfactory shots in 500 shot steps. Movement parameters have been set on “Walk on Spot”. Spectra were processed using FlexAnalysis software and the sophisticated numerical annotation procedure (‘SNAP’) peak detection algorithm, setting the signal-to-noise threshold at 5. Internal calibration was performed using the ^15^N ubiquitin peak ([M+H]+ average = 8669.5). Mass corresponding to ubiquitin (Ub initial = [M+H]+ average 8565.7), ubiquitin-adducts (Ub-K = [M+H]+ average 8735.7; Ub-T = [M+H]+ average 8709.6 m/z; Ub-S = [M+H]+ average 8695.8; Ub-glycerol = [M+H]+ average 8640.5 and Ub-Glucose = [M+H]+ average 8729.9) were added to the Mass Control List. Spectra were manually checked to ensure accuracy in calibration and peak integration. Peak areas were exported as .csv file using FlexAnalysis software and filtered using the previously described in-house GRID script ^29^. The percentage of discharge was calculated using the following equation:

### Recombinant expression of UBE2Q1 Wild type and mutants

Recombinant GST-fusion proteins were expressed in E. coli strain BL21 (DE3) cells. The cultures were grown in 2xTY or LB media containing 100 µg/ml ampicillin to an OD600 of 0.6-0.8 with 400 µM IPTG and were further allowed to shake overnight at 16°C. Cells were harvested in the following morning and were frozen and stored at -80. Cells were re-suspended in 50 mM Tris-HCl pH 7.5, 300 mM NaCl, 10% glycerol, 0.075% 2-mercaptoethanol, 1 mM AEBSF, and lysed by sonication. Bacterial lysates were clarified by centrifugation at 30,000 x g for 45 min and thereafter incubated with Glutathione Sepharose 4B resin for 45 minutes on low-speed rollers at 4 °C. The recombinant protein enriched resin was washed extensively first with a 50 mM Tris pH 7.5, 500 mM NaCl, and 10 mM DTT solution and then with the buffer containing physiological amount of salt (50 mM Tris-HCl pH 7.5, 150 mM NaCl, 10% glycerol, and 1 mM DTT). The GST tag was cleaved overnight on column incubation with 3C protease at 4°C. The purified proteins were dialysed into PBS buffer (pH 7.5 and 0.5 mM TCEP). Protein amount was determined using nanodrop while protein purity was established by SDS-page analysis. Proteins were flash frozen in liquid nitrogen and stored at -80 °C.

### Construction UBE2Q1∼Ub complex model

To construct a UBE2Q1∼Ub complex model we used UBE2D3∼Ub available crystal structures as a template (2QGX). The missing c-terminal residues in the crystal structure of UBE2Q1 minimal catalytic domain were traced using alphafold. The complex was finally obtained after several rounds of manual building in COOT further the model was refined using the webserver ^32^ to optimize the protein-protein interface.

### In vitro UBE2Q1 wild type or mutant autoubiquitylation assay

The autoubiquitylation assay included E2 (2.5 μM), UBE1 (0.5 μM), ATP (2 mM), MgCl2 (2 mM) and PBS pH 7.5. The reaction was started by adding the UBE2Q 1 full length or minimal catalytic domain and or UBE2Q1 mutants and incubating the mixture at 30°C for 30 minutes. The reaction was stopped using 4XLDS buffer at the indicated time point and further visualised using the SDS-PAGE. The images were captured using Chemidoc.

### In vitro Β4GalT1 peptide ubiquitylation assay

The in vitro Β4GalT1 peptide ubiquitylation assay were performed as described above for autoubiquitylation assay with the addition of Β4GalT1 peptide. The reaction mixtures were incubated at 30 degrees for 1 hour. The reaction was stopped using 4XLDS buffer at the indicated time point and further visualised using the SDS-PAGE. The images were captured using Chemidoc.

### Sourcing and analysis of E2 conjugating enzymes expression profiling in human tissues

Raw data from “A Quantitative Proteome Map of the Human Body” ^33^ were downloaded from Proteome Xchange (PXD016999) ^34, 35^ and searched against Uniprot SwissProt Human containing isoforms (downloaded on 05 October 2021) using MaxQuant (v2.0.3.1)^36^. MS1 intensities per channel were estimated by weighting the MS1 intensity with the TMT intensities. Protein copy numbers were estimated using the proteomics ruler plugin ^37^in Perseus (v2.0.3.0)^38^. Data were further analysed and plotted using Python (v3.9.0) and the packages Pandas (v1.3.3), Numpy (v1.19.0) and Plotly (v5.8.2).

### Animals and Tissue processing for immunoblotting analysis

The C57BL/6J mice were obtained from Charles River Laboratories (Kent-UK) and housed in a specific pathogen–free facility in temperature-controlled rooms at 21°C, with 45 to 65% relative humidity and 12-hour light/12-hour dark cycles with free access to food and water and regularly monitored by the School of Life Science Animal Unit Staff. All animal studies were approved by the University of Dundee Ethical Review Committee and performed under a U.K. Home Officer project license. Experiments were conducted in accordance with the Animal Scientific Procedures Act (1986) and with the Directive 2010/63/EU of the European Parliament and of the Council on the protection of animals used for scientific purposes (2010, no. 63).

6-month-old C57BL/6J mice were killed by cervical dislocation and peripheral tissues were rapidly washed in ice-cold phosphate-buffered saline (PBS) and snap frozen in liquid nitrogen.

Whole brain was dissected out from the skull, rapidly washed in ice-cold PBS and placed on an ice- cooling plate under a stereomicroscope for brain sub-regions microdissection. Brain sub-regions, such as olfactory bulbs, cortex, hippocampus, striatum, hypothalamus, thalamus, midbrain, cerebellum, brainstem and spinal cord were dissected and collected in a single 1.5 ml microcentrifuge tube and snap-frozen in liquid nitrogen. Tissue samples were stored at −80°C until ready for processing. All tissues were weighed and homogenised in 5X volume for mg of tissue of ice-cold lysis buffer containing: 50 mM Tris/HCl pH 7.5, 1 mM EDTA pH 8.0, 1 mM EGTA pH 8.0, 1% Triton X-100, 0.25 M sucrose, 1 mM sodium orthovanadate, 50 mM NaF, 10 mM sodium glycerol phosphate, 10 mM sodium pyrophosphate, 200 mM 2-chloroacetamide, phosphatase inhibitor cocktail 3 (Sigma- Aldrich) and complete protease inhibitor cocktail (Roche). Tissue homogenization was performed using a probe sonicator at 4°C (Branson Instruments), with 10% amplitude and 2 cycles sonication (10 seconds on, 10 seconds off). Crude lysates were incubated at 4°C for 30 min on ice, before clarification by centrifugation at 20,800 x g in an Eppendorf 5417R centrifuge for 30 min at 4°C. Supernatants were collected and protein concentration was determined using the Bradford kit (Pierce).

## Results

### Discovery of new non-canonical E2 conjugating enzymes

Because of the recent discovery of the unexpected ability of E3 ligases to ubiquitylate non- canonical residues, we asked whether other enzyme within the ubiquitin cascade, particularly E2 conjugating enzymes, could also been intrinsically reactive toward non-canonical residues. We therefore developed a MALDI-TOF MS based assay to detect the formation of ubiquitin adducts resulting from E2 conjugating discharge activity of ubiquitin on different nucleophiles (Fig. 1a). The E2 Discharge MALDI-TOF assay relies on detection of the ubiquitin adduct formed in presence of a nucleophile on which the E2s will discharge ubiquitin (Fig. 1a and b). The ubiquitin adducts can be directly detected as a consequence of E2 activity over time and absolute and relative quantification is assessed through the use of an internal standard (^15^N ubiquitin) (Fig. 1b). A panel of 23 recombinantly-expressed E2 conjugating enzymes (2.5 µM final, see Table 1) was tested for their ability to discharge ubiquitin onto Ac-lysine (Ac-K), Ac-threonine (Ac-T), Ac-serine (Ac-S), glycerol and glucose. Reactions were conducted at 30°C and incubated for 1 h in presence of the indicated nucleophiles (50 mM final). E2s known to work with RING-type E3s have E3- independent reactivity towards lysine. The majority of E2 conjugating enzymes discharged ubiquitin in presence of Ac-lysine while no corresponding Ub-adduct was observed in presence of either Ac-serine, Ac-threonine, glycerol or glucose (Fig. 1c). Consistent with previous literature, the HECT specific E2 conjugating enzyme, UBE2L3, did not discharge on lysine ^39^. Also, Ube2W exhibits no intrinsic activity towards free lysine as previously reported ^40, 41^ since this particular E2 specifically attaches ubiquitin to the N-terminal α-amino group of proteins ^42^. The UBE2J2 conjugating enzyme has been previously reported to be intrinsically reactive toward lysine but, unexpectedly, has also been found to be reactive toward serine ^43^. In accordance with previous studies ^43^, our data show that UBE2J2 is able to ubiquitylate glycerol, glucose, serine and lysine but – interestingly - not threonine, indicating that a hydroxyl group alone was not sufficient to confer UBE2J2 reactivity toward its substrate. Strikingly, two E2s, UBE2Q1 and UBE2Q2, were able to conjugate ubiquitin to serine, threonine, glycerol and glucose residues but showed relatively low reactivity toward lysine residues (Fig. 1c). Interestingly, while both UBE2Q1 and UBE2J2 were able to ubiquitylate the more complex sugar maltoheptaose (See Sup. Fig.1a and b), UBE2Q1 did so more efficiently than UBE2J2 (See Sup. Fig.1c). To further confirm and characterize the ability of UBE2D3, UBE2J2, UBE2Q1 and UBE2Q2 to ubiquitylate hydroxylated substrates, we tested them for discharge activity over time (Fig. 1d). UBE2D3 showed lysine-specific discharge throughout the time course experiment. UBE2Q1 showed discharge activity on all three nucleophiles but with an higher activity rate toward Ac-T compared to Ac-S and Ac-K, while UBE2Q2 showed similar reactivity toward Ac-S and Ac-T and reduced discharge on Ac-K. UBE2J2 actively discharged on both lysine and serine residues with similar rates while it showed no discharge on threonine throughout the time course experiment.

**Figure 1.**
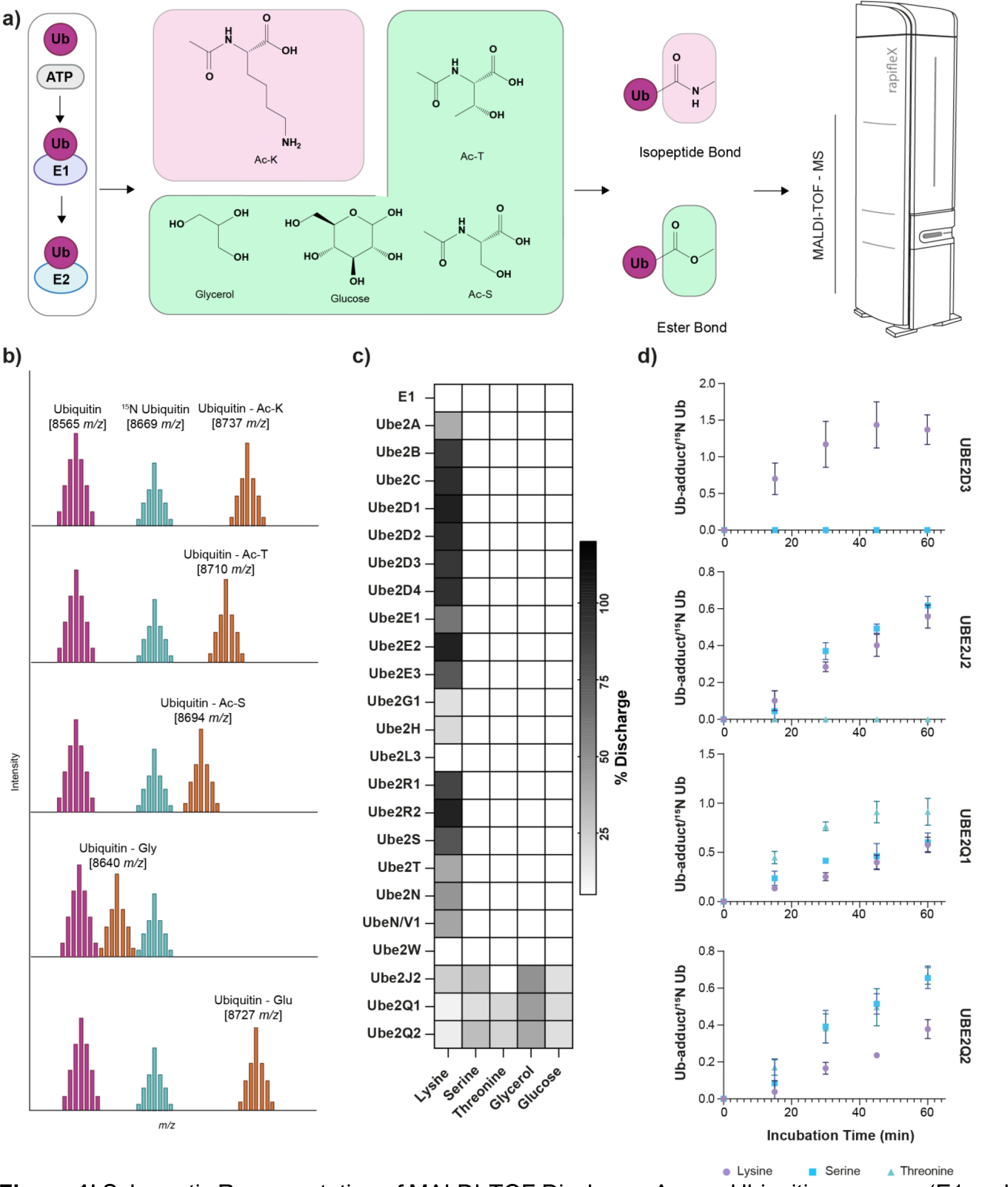
Schematic Representation of MALDI-TOF Discharge Assay. Ubiquitin enzymes (E1 and E2) are incubated with ATP/MgCl2 solution and different nucleophiles. Samples are then analysed by MALDI-TOF MS. Relative quantification of E2 discharge activity is obtained by use of an internal standard (^15^N Ubiquitin) (panel b). 23 E2 conjugating enzymes were tested for their discharge ability (c). Canonical and Non-canonical discharge of UBE2D3, UBE2Q1, UBE2Q2 and UBE2J2 was further validated in a time course experiment (d)

### UBE2Q1 auto ubiquitylates on non-lysine residues

UBE2Q1 undergoes extensive auto ubiquitylation *in vitro* (Fig. 2a, lane 2). To test the chemical nature of the bond that originated UBE2Q1 autoubiquitylation bands, the sample pH was either reduced with Sodium Hydroxide (Fig. 2a, lane 3), treated with β-mercaptoethanol (βME) to specifically cleave thioester bond (Fig. 2a, lane 4) or with hydroxylamine to cleave both ester and thioester bonds, (Fig. 2a, lane 5). The sensitivity of UBE2Q1 autoubiquitylation smear to mild alkaline and hydroxylamine treatment but not thiol reduction with β-mercaptoethanol indicated that such auto-modification results from the formation of ester rather than isopeptide or thioester bond. Several UBE2Q1 autoubiquitylation bands also disappeared in presence of the deubiquitinating enzyme (DUB) JOSD1, a member of the Machado-Josephin disease DUB family previously reported to specifically cleave the ester bond linking ubiquitin to threonine substrate but unable to hydrolyse the isopeptide bond linking ubiquitin to lysine ^44^ (Fig. 2a, lane 6). JOSD1 treatment was coupled with β-mercaptoethanol reduction (Fig. 2a, lane 7): no difference was observed compared to the JOSD1 treatment alone, further confirming that JOSD1 mediated cleavage is restricted to ester-bond conjugated ubiquitin. USP2, a DUB able to cleave both ester and isopeptide bond, removed all UBE2Q1 autoubiquitylation bands.

**Figure 2.**
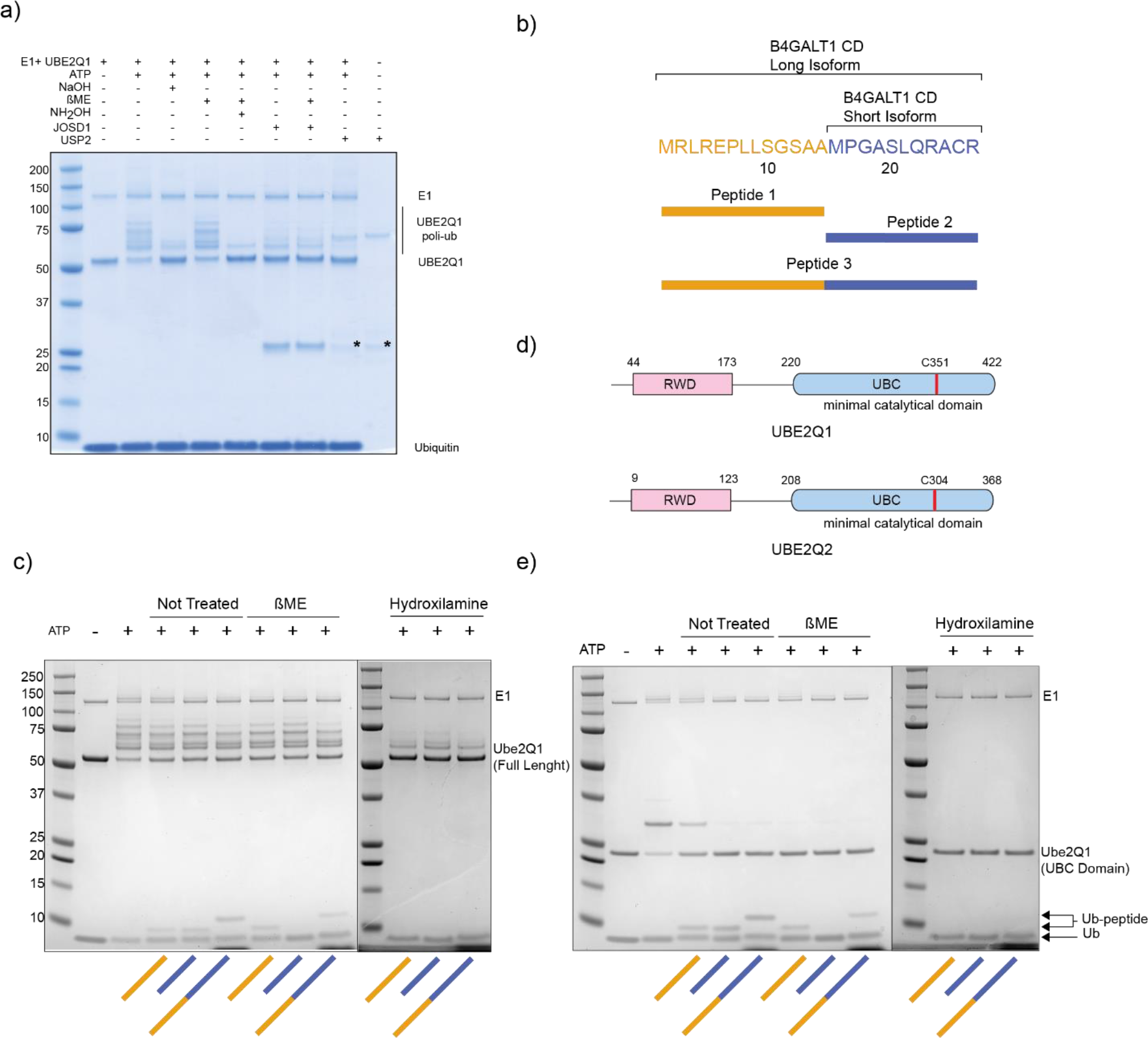
UBE2Q1 autoubiquitylation bands are sensitive to hydroxylamine and sodium Hydroxide treatment (a). Schematic of B4GALT1 cytoplasmic domain and peptides synthesized in this study (b). Full length UBE2Q1 directly ubiquitylates B4GALT1 on cysteine and serine residues (c). UBE2Q1 and UBE2Q2 schematic representation (d). UBE2Q1 UBC-domain is sufficient to directly ubiquitylate B4GALT1 domain.

### UBE2Q1 directly ubiquitylates B4GALT1 cytoplasmic domain

UBE2Q1 substrates are currently unknown, however UBE2Q1 was found to directly interact with the cytoplasmic domain (CD) of the Golgi resident Beta-1,4-galactosyltransferase 1 (B4GALT1) ^45^. The B4GALT1 gene encodes two isoforms that differ for the length of the CD, a short segment that encodes 24 amino acids at the protein N-termini. The full-length isoform includes the distal component of the B4GALT1 CD domain (13 amino acids) while the short isoform encodes only its proximal portion (11 amino acids). Strikingly, neither B4GALT1 shorter isoforms contain a lysine residue, but both of them possess 3 serine and one cysteine. We hypothesized that UBE2Q1 ubiquitylates B4GALT1 CD on these non-canonical residues. Peptides belonging to the long isoform (peptide 1), short isoform (peptide 2) and the full length B4GALT1 CD (peptide 3) were synthesized and incubated with E1 activating enzyme, UBE2Q1 and ATP/MgCl2. UBE2Q1 directly ubiquitylated both peptide 1 and 3 on a serine residue, demonstrated by the sensitivity to hydroxylamine treatment but not to β - mercaptoethanol (Fig. 2c). The ubiquitylation of peptide 2 was instead mediated by thioester bond with the cysteine, as the sensitivity to β – mercaptoethanol indicates.

UBE2Q1 and UBE2Q2 are characterized by an extended N-terminus, that includes a protein domain (RWD) shared by RING finger-containing proteins, WD-repeat-containing proteins, and yeast DEAD (DEXD)-like helicases ^46^ (Fig. 2d). RWD domains have been suggested to be substrate recognition domains for ubiquitin-conjugating enzymes ^47^ but their specific function is currently not completely understood. We speculated that the extended N-terminus of UBE2Q1 might play a role in the interaction and the recognition of the B4GALT1 CD. We therefore tested a UBE2Q1 construct - containing only the UBC fold domain (UBE2Q1 UBC domain) - for its ability to directly ubiquitylate the B4GALT1 CD. UBE2Q1 UBC domain was still efficiently ubiquitylating the B4GALT1 CD on serine residues therefore suggesting that this domain is sufficient to recognize the B4GALT1 CD sequence *in vitro* in absence of its N-terminus or a cognate E3 ligase (Fig. 2e). Notably, UBE2Q1 UBC domain did not undergo extensive autoubiquitylation (Fig. 2e), suggesting that the autoubiquitylation events produced by full length enzyme are confined within its extended N-terminus.

### UBE2Q1 uses a non-canonical catalytic triad for substrate ubiquitylation

Canonical E2s are characterized by a conserved Histidine Proline/Cysteine- Asparagine (HP/CN) motif in the active site ^48^. Notably, the UBE2Q family and UBE2J2 lack these conserved catalytic residues that are fundamental for the reactivity toward lysine and the formation of the isopeptide bonds ^49^ (Sup. Fig. 2). We therefore asked which residues within UBE2Q1 active site played a role for the activity of this class of enzymes.

To understand the underlying mechanism allowing UBE2Q1 to both interact with and discharge ubiquitin onto the substrate, we constructed an UBE2Q1-Ub model using as a scaffold template the available structure of UBE2D3 – a canonical E2 conjugating enzyme – loaded with ubiquitin (UBE2D3∼Ub) and used Alpha fold and the COOT for modelling the UBE2Q1-ubiquitin interaction. The structural comparison with the UBE2D3∼Ub complex highlighted a substantially different mode of interaction between ubiquitin C-terminus and the respective E2s. In the UBE2D3∼Ub complex, the thioester bonded ubiquitin barely interacts with the residues in the proximity of the catalytic cysteine, unlike in the UBE2Q1∼Ub modelled complex where the C- terminal of the ubiquitin is deeply buried within the UBE2Q1 active site (Fig 3 a). A closer view of the UBE2Q1∼Ub modelled complex revealed that the interactions between the UBE2Q1 and the ubiquitin consist mainly of hydrogen bonds and hydrophobic interactions. Residues present at the interface between ubiquitin and the UBE2Q1 UBC fold (Fig. 3a and Sup. Fig. 3) and C-terminus were systematically mutated and tested for their ability to impact either the ubiquitin loading (loading-defective mutants) or the ubiquitin discharge onto B4GALT1 (discharge-defective mutants). All mutants (Fig. 3 b-d and Sup. Fig 3 a-c) were tested by MALDI-TOF discharge assay and ubiquitylation of B4GALT1 peptide 1 (Fig. 3 c and d). Three residues, Y343, H409 and W414 were identified as critical for the ability of UBE2Q1 to discharge on both canonical and non- canonical residues while leaving unaffected the ubiquitin loading step (Fig. 3d). Swapping histidine 409 with asparagine did not rescue UBE2Q1 enzymatic activity, therefore suggesting that histidine mediates essential hydrophobic interaction and/or necessary hydrogen bonds with the substrate. Similarly, mutating W414 with either phenylalanine or with glutamine did not rescue UBE2Q1 activity, thus highlighting the specific role that tryptophan 414 plays either in promoting the catalysis of the ester bond and/or in the recognition and binding of the substrate. These results indicate that UBE2Q1 uses an alternative catalytic triad, comprised of a non-sequential YHW motif (See Sup. Fig. 2), to recognize non-canonical substrates and to drive the formation of ester bonds.

**Figure 3.**
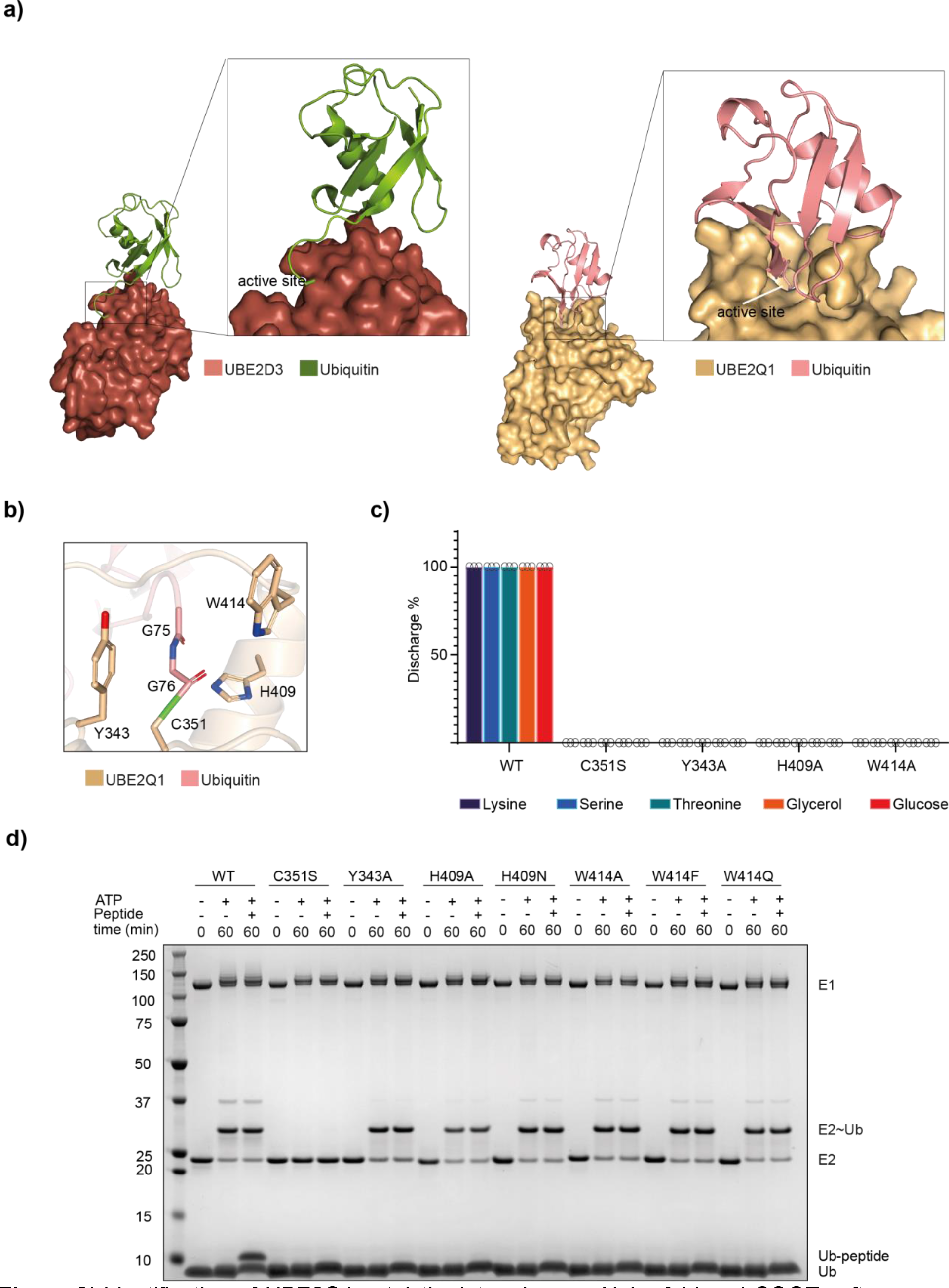
Identification of UBE2Q1 catalytic determinants. Alpha fold and COOT software were used to model the interaction between UBE2Q1 and ubiquitin using the UBE2D3-Ub available structure as reference (panel a). Model of Y343, W414 and H409 interacting with ubiquitin c-terminus (b). Indicated UBE2Q1 mutants were tested for their ability to discharge on the indicated nucleophiles by MALDI-TOF MS (c) and for the ubiquitylation of B4GALT1 peptide 1 (d).

### UBE2Q1 prefers threonine over serine

The initial MALDI-TOF E2 discharge assay time course dataset (Fig. 1d) was suggestive of an underlying preference of UBE2Q1 toward threonine rather than serine or – even more markedly - lysine. The.B4GALT1 CD is highly evolutionary conserved in mammals. Serine 11 and 18 are retained or conservatively mutated in all the analysed mammalian sequences, while serine 9 is present only in primates (Fig. 4a). To test the preference of UBE2Q1 toward serine, threonine or lysine, the two serines present within B4GALT1 peptide 1 were systematically mutated into threonine, lysine or alanine. Substituting serine 9 in peptide 1 with alanine reduced the amount of ubiquitylation of the peptide by around 50%, suggesting that the ubiquitylation event is distributed among serine 9 and 11. Mutating serine 9 into alanine and serine 11 lysine completely (and vice versa) abolished peptide ubiquitylation, further confirming an intrinsic preference of UBE2Q1 toward residues with a hydroxyl group (see Fig. 4b). Interestingly, UBE2Q1 showed a marked increase in the ubiquitylation band when serine 11 and serine 18 were mutated into threonine but not when the S>T modification was inserted in position 9 of peptide 1 (See Fig. 4 b and c). Remarkably, the substitution of serine 18 into threonine in peptide 2 led to a β – mercaptoethanol resistant ubiquitylation band, suggesting that UBE2Q1 has a strong preference for threonine even in the presence of the thiol scavenging cysteine residues. The result indicated that UBE2Q1 prefers threonine over cysteine, over serine as substrate.

**Figure 4.**
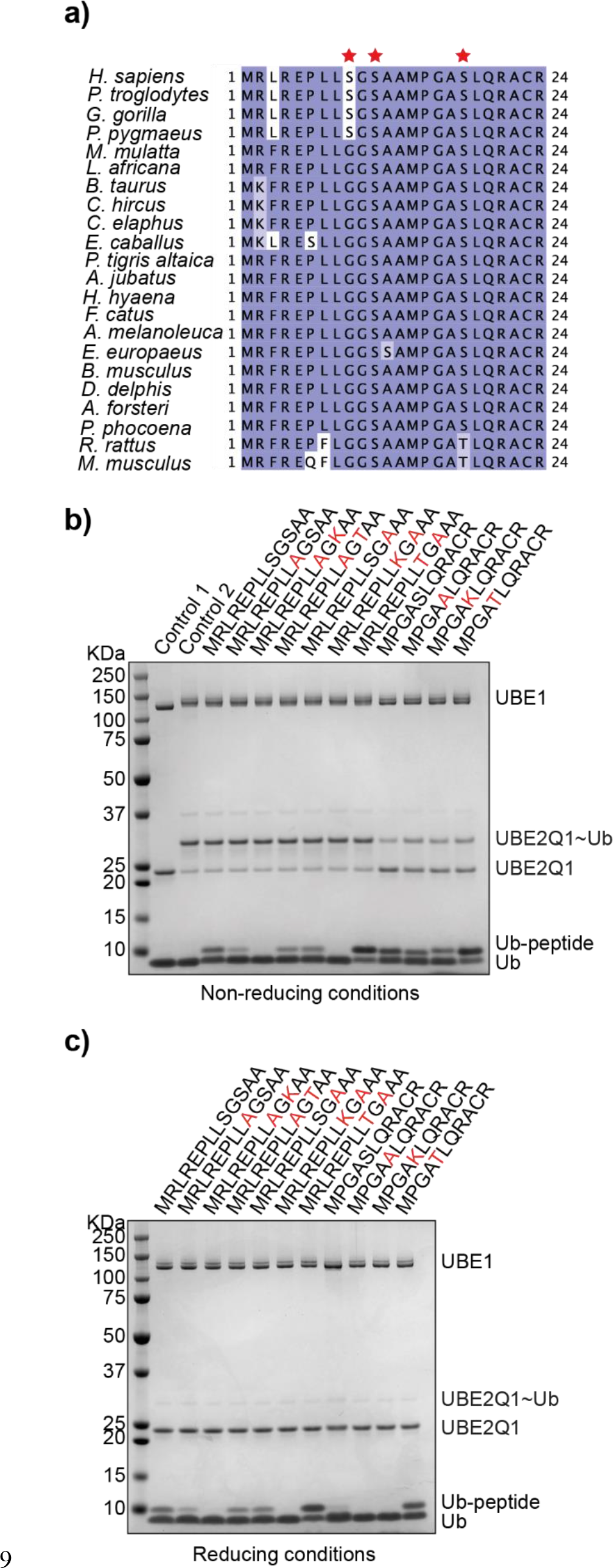
B4GALT1 Cytoplasmic Domain (CD) is highly conserved. Sequence alignment of B4GALT1 CD domain in mammals (a). Serine residues were systematically mutated into Alanine, Lysine or Threonine and tested for UBE2Q1 mono-ubiquitylation in non-reducing (b) and reducing conditions (c).

### UBE2Q1 is highly expressed in the brain

While the role of UBE2J2 in ERAD has been characterized, the biological role(s) of the UBE2Qs family is unclear. To determine whether UBE2Qs are tissue-specific or tissue-enriched we interrogated a publicly available high quality proteomic dataset in which 32 human tissues were analysed using quantitative proteomics^33^. Interestingly, of about 40 E2s encoded in the human genome, only 21 were expressed in sufficient quantities to be detected in the dataset (Fig. 5a). UBE2Q1 and UBE2J1 were identified in all analysed tissues, while UBE2Q2 and UBE2J2 were not detected, therefore suggesting that these E2 might be either relatively low abundant or expressed in other tissues or under specific biological conditions. Interestingly, UBE2Q1 was found to be relatively more expressed in brain and in testis (see Fig.5) suggesting a specific role in these tissues. We developed an in house UBE2Q1 antibody to specifically detect and verify the expression of UBE2Q1 in different cell lines and tissues. Sections of mice brain and other tissue (liver, spleen, kidney and heart) were collected from 4 different mice and tested for UBE2Q1 expression levels. Indeed, UBE2Q1 was found to be highly expressed in all brain regions; a lower expression was observed in the spleen, liver and kidney while it was detected in the heart (Fig. 5c).

**Figure 5.**
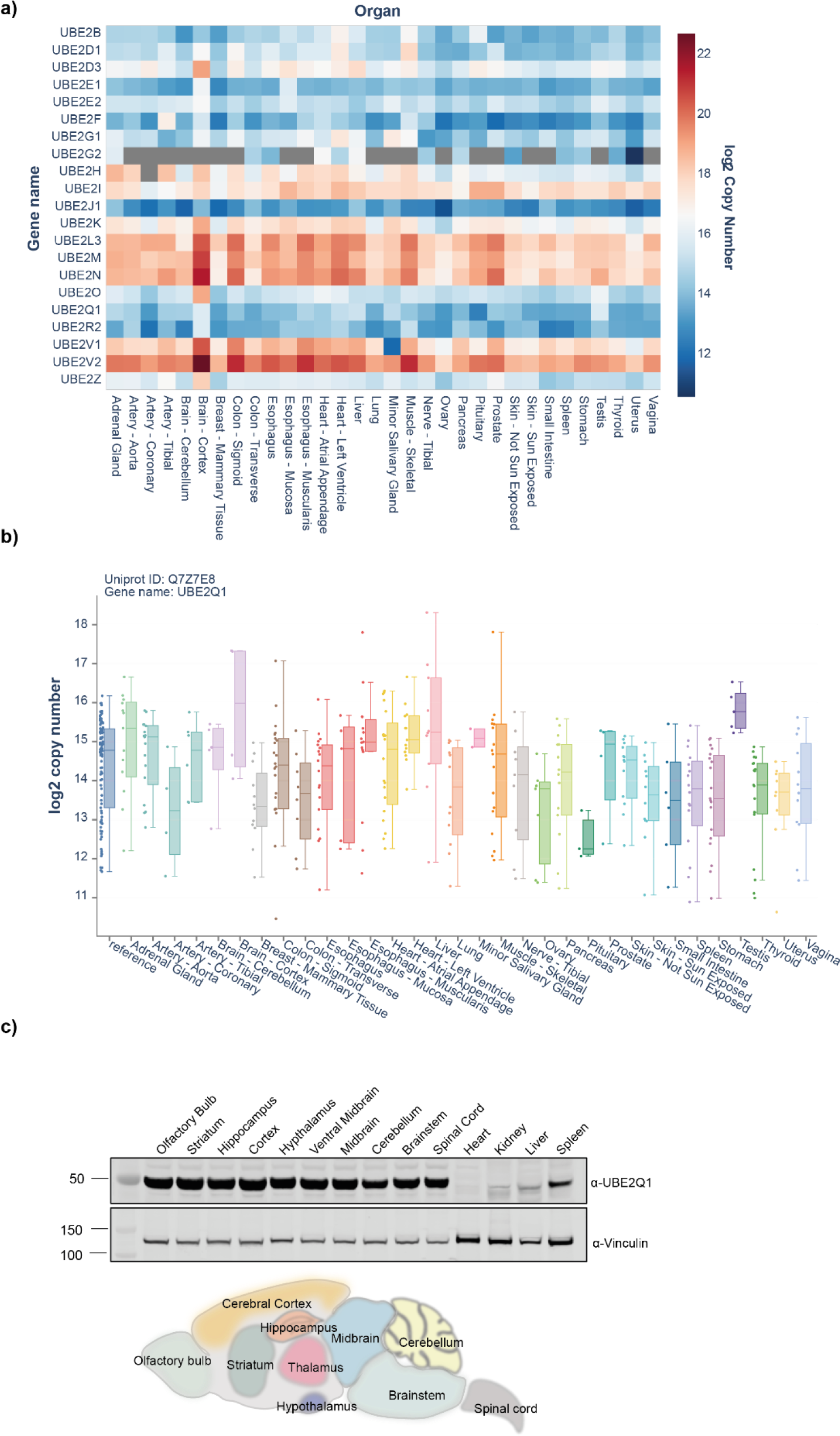
Relative expression of E2 conjugating enzymes in human and mouse tissues. E2 in vivo expression profiling obtained by data mining of publicly available quantitative proteomic dataset of 32 human tissues (a). Relative expression of UBE2Q1 in indicated human tissues (b). Western blot validation of UBE2Q1 expression in the indicated mouse tissues (c).

### Preference of serine and cysteine ubiquitylation over lysine by UBE2J2 is independent of ligase interactions

To date, the intrinsic ability of the Ube2J2∼Ub conjugate to react with serine or threonine has not been directly demonstrated, so neither the structural nor chemical determinants for hydroxyl attachment of ubiquitin have been identified. Both UBE2J2 and B4GALT1 are localized within secretory pathway and are membrane bound. We therefore speculated that also UBE2J2 might also ubiquitylate B4GALT1 CD.

Indeed, UBE2J2 efficiently ubiquitylated B4GALT1 peptide 1(Fig. 6a lane 3); however, mutating serine on position 9 to alanine substantially reduced the discharge of the ubiquitin onto the peptide (Fig 6a Lane 4). On the other hand, serine mutation on position 11 to alanine showed decreased ubiquitylation compared to the wild type, but discharge did not cease (Fig 6a lane 7). This differential effect on the pattern of the ubiquitylation highlights the specificity of the serine position on the peptide. To test the preference of serine over lysine, we mutated both serines at position 9 and 11 to lysine within B4GALT1 peptide 1 and observed little or no ubiquitylation (Fig 6a lanes 5 and 8). Replacing both serines with threonine in either position did not rescue the pattern of ubiquitylation, as observed in peptide 1 (Fig 6a lanes 6 and 9). Thus, showing that serine is preferred not only over lysine but over threonine as well. Interestingly, peptide 2 – the cysteine containing portion of B4GALT1 CD – was strongly ubiquitylated by UBE2J2 (Fig. 6a lanes 10-13). The nature of peptide 2 ubiquitylation was thioester based as demonstrated by the sensitivity of these adducts to β-mercaptoethanol treatment (Sup. Fig 4a lanes 10-13). Also, the appearance of UBE2J2 autoubiquitylation bands that are sensitive to β-mercaptoethanol (by comparing Fig. 6a with Sup. Fig. 4a) indicates an intrinsic reactivity of UBE2J2 toward cysteine residues. Overall, these results suggest that UBE2J2 strongly favors cysteine over serine, while no lysine ubiquitylation is detected within the B4GALT1 CD.

**Figure 6.**
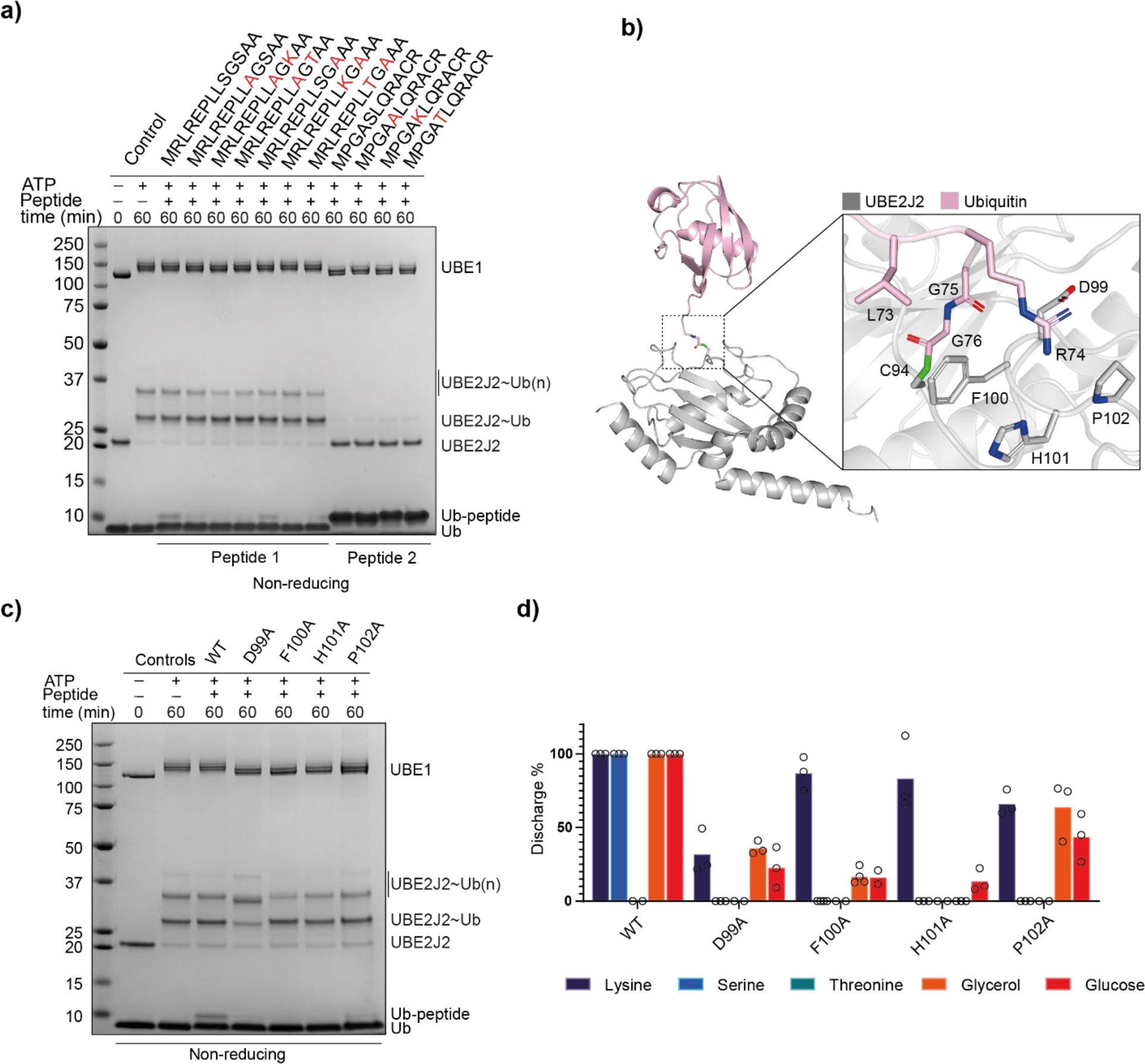
Identification of UBE2J2 catalytic determinants. UBE2J2 ubiquitylates B4GALT1 CD on peptide 1 and peptide 2 (a). Predicted conformations of the interaction between UBE2J2 and ubiquitin (b). UBE2J2 active site and residues predicted to be relevant for enzymatic activity (c). UBE2J2 mutants tested for B4GALT1 CD peptide 1 ubiquitylation (d) and for discharge on the indicated nucleophiles by MALDI-TOF MS.

Interestingly, since all these discharge assays were done in the absence of an E3 ligase, this further established that the ability to discriminate between residue side chains is independent of any E3 ligase.

### Residues in the vicinity of catalytic cysteine are critical for ubiquitin discharge

To characterise the molecular interactions that govern the assembly of the UBE2J2∼ub complex in the absence of any crystal structure, we built an UBE2J2∼ub model using the alpha fold and COOT. Unlike UBE2Q1∼Ub, the UBE2J2∼Ub model shows multiple conformations possible for ubiquitin bound to the catalytic cysteine (Fig. 6b). This flexible ubiquitin may have the potential to interact with the residues in the vicinity of the catalytic cysteine. Interestingly, sequential residues interacting with ubiquitin are found to be disordered in the UBE2J2 apo crystal structure reported (PDB ID: 2F4W). This stretch is highly mobile and might attain ordered conformation when an incoming ubiquitin forms a thioester bond with the catalytic cysteine (Sup. Fig. 5). The complex after energy minimisation to remove short contacts was assessed to identify the critical residues that may play a role in the discharge onto the substrate. Some of the sequential residues found interacting with the C-terminal of the ubiquitin bound to the catalytic cysteine are D99, Y100, H101, P102 and D103 from various conformations. The UBE2J2 also interacts through L129 with the bound ubiquitin (Fig. 6b inset). To functionally validate the model, we mutated the highlighted residues to alanine and observed the pattern of ubiquitylation discharge onto peptide 1. None of the mutants showed significant ubiquitin loading defects, however they could no longer discharge ubiquitin onto peptide 1 (Fig. 6c), which showed accurate prediction in the identification of residues relevant for stabilization of the ubiquitin C-terminus and substrate ubiquitylation. We further assessed these mutations for their ability to impact the discharge onto nucleophiles by MALDI-TOF MS (Fig. 6d). Mutating F100 and H100 into alanine nearly completely abolished discharge on hydroxyl group containing molecules - serine, glycerol and glucose but only partially reduced the discharge on lysine, therefore suggesting that these are residues relevant for the catalysis of ester bond. P102A showed a mixed phenotype, with about 50% reduction of the lysine-mediated discharge and 70% and 85% reduction in the reactivity toward glycerol and glucose respectively. On the other hand, the D99A mutant showed a 50-70% reduction in the formation of Ubiquitin-K adducts. Notably, the UBE2J2 D99 residue aligns with the D87 residue in the canonical UBE2D3 E2 conjugating enzyme (See Sup. Fig. 6): this residue was previously identified for having a general role in lysine reactivity^39^. Overall, these results define UBE2J2 as an hybrid E2 conjugating enzyme: similarly to canonical E2s, UBE2J2 possesses a sequential histidine and proline residues that are highly conserved and structurally necessary in canonical E2s although dispensable for isopeptide bond formation ^50^. However, UBE2J2 lacks the asparagine residue, previously deemed essential in the isopeptide bond catalysis ^50^ while retaining a critical aspartic acid (D99) in line with other – lysine specific - E2s ^39^. Despite sharing many similarities with canonical E2s, UBE2J2 possesses an intrinsic and E3 independent reactivity toward serine and cysteine that relies on multiple sequential residues within the active site.

## Discussion

E2 conjugating enzymes play an upstream role within the ubiquitylation cascade: while E3 ligases confer substrate specificity, E2s dictate the catalytic activity that leads to the attachment of ubiquitin to the substrate. By interacting with multiple RING E3 ligases, E2s have the potential to tag a wide range of substrates with ubiquitin. The majority of E2s have been reported to possess intrinsic reactivity toward lysine, this being assessed through SDS-page based assays that rely on the visualization of bands corresponding to the E2 enzyme loaded with ubiquitin and its disappearance in presence of high concentration of a nucleophile. While extensively used, the SDS-page based assay presents some limitations related to its intrinsic low resolution, including the impossibility to resolve adducts formed with commonly used buffer reagents such as glycerol or sucrose – and the impossibility to discriminate between bands that correspond to the E2 being loaded with ubiquitin or autoubiquitylation events. In 2018, we develop a MALDI-TOF MS-based assay that allows the direct quantification of E2 and E3 activities based on the disappearance of free ubiquitin in presence of productive E2/E3 pairs ^28^. The use of MALDI-TOF mass spectrometry allowed for the detection of unexpected ubiquitin – glycerol adduct as results of the UBE2Q1 and UBE2Q2 non-canonical activity. UBE2Q1 and UBE2Q2 were subsequently found to actively discharge onto several hydroxyl containing molecules, which sets them apart from canonical E2s.

UBE2J1 and UBE2J2 were previously identified and named as Non-Canonical Ubiquitin-Conjugating Enzyme – NCUBE1 and NCUB2 because of the lack of the sequential Histidine-Proline-Asparagine (HPN) motif, that is highly conserved in mostly known and canonical E2s ^48, 49^. The function of the HPN motif is thought to be both structural and functional: histidine and proline are structurally important in forming the E2 active site^50^, while the asparagine residue is important for mediating the catalysis of an isopeptide bond between ubiquitin and a substrate lysine ^50^. Besides the HPN triad, two aspartic acid residues were also identified as relevant for their hydrogen-bond based interaction with the conjugated ubiquitin ^39^. Our structural motif scanning and functional assay validation led to the identification of critical - non-sequential - residues in the UBE2Q1-Ub model responsible for the stability of ubiquitin C-terminal tail and the catalysis of both ester and isopeptide bonds.

Mutating UBE2Q1 Histidine 409 into alanine abolished the discharge activity onto all tested nucleophiles, demonstrating that this residue is well conserved and necessary for catalysis of both canonical and non-canonical substrates. We further identified two highly hydrophobic residues within the UBE2Q1 catalytic site, tryptophan 414 and phenylalanine 343, that contribute to the formation of a hydrophobic pocket essential for the interaction between the ubiquitin C-terminus and the serine and threonine residues present in the substrate.

We also found that UBE2J2 possesses a rather peculiar catalytic triad, where histidine 101 is essential for the formation of the ester bond while dispensable for the discharge on lysine and the catalysis of isopeptide bond. Instead, UBE2J2 relies on an aspartic acid residue (D99), located upstream of the Histidine-Proline sequence, to actively discharge on lysine residues. These results indicate that UBE2J2 uses different catalytic residues to actively interact with different nucleophiles. UBE2J2 catalytic site appears therefore to be an E2 “hybrid”, featuring residues belonging to both canonical E2s - a Histidine and Proline sequential motif and a key catalytic aspartic acid residue – and the ability to ubiquitylate lysine but also cysteine, serine and complex sugars.

Besides their communality as non-canonical E2s, UBE2Qs and UBE2J2 have also several dissimilarities. UBE2Q1 strongly prefers threonine vs serine, while UBE2J2 does not ubiquitylate threonine at all. All E2s enzymes are intrinsically reactive toward thiols as requisite for accepting the thioester-linked ubiquitin from the E1 activating enzyme. Nevertheless, UBE2J2 showed a remarkable reactivity toward the cysteine-containing portion of B4GALT1 CD compared to UBE2Q1. It might be argued that the *in vitro* scavenging activity of the cysteine residue within peptide 2 does not translate into a genuine *in vivo* preference. However, UBE2J2 was also previously reported as the E2 responsible for the ubiquitination of the MHC I intracytoplasmic tail on a cysteine when paired to the viral RING E3 ligases MIR1 and MIR2^20, 22^. Moreover, UBE2J2 autoubiquitylation profile is supportive of an intrinsic preference of UBE2J2 toward cysteine, we therefore propose that UBE2J2 is a cysteine and serine specific E2 conjugating enzyme.

Normally E2∼Ub conjugates present themselves in a “open” conformation with low rates of ubiquitin transfer in the absence of an E3 ligase to avoid cycles of conjugation and off-target ubiquitylation. RING E3s bring the substrate and the E2∼Ub conjugate together and stabilize the E2∼Ub conjugate in the active “closed” conformation required for lysine ubiquitylation ^51–54^. E2s that abide by this model do not directly ubiquitylate a substrate in absence of their cognate E3. The exception to this rule is represented by UBE2I/Ubc9, a sumo-specific E2 conjugating enzyme, which, in the absence of E3, can directly sumoylate a target lysine embedded within a consensus motif ψK*X*(E/D) (ψ indicates a hydrophobic amino acid, whereas *X* indicates any amino acid) ^55^. Similarly, both UBE2J2 and UBE2Q1 were found able to directly ubiquitylate the 24 amino acid B4GALT1 cytoplasmic domain also in absence of a RING E3 ligase. In the case of UBE2Q1, the E3-independent ubiquitylation was not mediated by the extended N-terminus, therefore suggesting that some other interactions between the C-terminus ubiquitin and the UBE2Q1 catalytic domain allow for the direct binding to the polypeptide. The reactivity of both UBE2Q1 and UBE2J2 toward the B4GALT1 cytoplasmic domain suggest that these E2 adopt and intrinsically more reactive conformation even in absence of a cognate E3 ligase or that their UBC domain is posed to recognize a specific short sequence within their substrates.

Interestingly, ubiquitylation of a lysine-free, short cytoplasmic domain belonging to membrane proteins is not uncommon. The cytoplasmic tail of T-cell receptor α, consisting of the residues RLWSS, was previously found to be ubiquitylated by the combined action of UBE2J2 and HRD1 ^23^. In this case, the exact position of the serines within the tail is not as important as is the nature of the surrounding residues, where less hydrophobic residues enhance ubiquitylation on serine. UBE2J2 was also found to ubiquitylate the cytoplasmic tail of the major histocompatibility complex class I heavy chains by interacting with the γ-HV68 murine virus K3 ligase (mK3) ^43^. Similarly, the viral protein VPU can ubiquitylate the cytoplasmic tail of CD4^56^ and the cytoplasmic domain of BST-2/Tetherin^57^ on serine and threonine residues. All of these membrane proteins only have few residues that are accessible for ubiquitylation. It might be speculated that non-canonical E2- conjugating enzymes have evolved to target those short, lysineless sequences. UBE2Qs and UBE2Js are not located on the same cellular compartment. UBE2J2 is bound to the membrane of the endoplasmic reticulum, where it is required for ubiquitination of multiple ER-associated protein degradation (ERAD) substrates^43^. By contrast, UBE2Q1 is reported as located mainly in the cytoplasm. The difference in cellular compartment location of these E2s might suggest that these E2s affect different ubiquitylation substrates within different cellular compartments. The high expression of UBE2Q1 in the brain correlates with its reported role in traumatic brain injury and frontotemporal dementia ^58, 59^. UBE2Q1 has also being found to have a pleiotropic effect on fertility by playing a fundamental role during the implantation and development of embryos and subsequent pregnancy viability ^60^. Indeed, UBE2Q1 -/- female mouse shows significantly reduced fertility rates. We anticipate that the discovery of UBE2Q1 non-canonical activity will help to fully resolve the molecular mechanisms that drive such dramatic phenotype.

Notably, a third member of the UB2Q family, UB2QL1, known to be important in the clearance of damaged lysosomes ^61^, was inactive in our in vitro assay. However, it is likely to possess the same non-canonical activity of the other UB2Q1 family members. Similarly, UB2J1 was also found inactive *in vitro*. This suggests that both enzymes might require the cellular environment, specific co-factors or specific posttranslational modifications to function. In total, 5 out of about 39 E2 enzymes are likely to possess non-canonical activity. In summary, the growing number of ubiquitin enzymes able to target amino acids other that lysine, particularly E2s, which act upstream of E3 ligases, suggest that there is a vast pool of potential substrates that might be subjected to non-canonical ubiquitin regulation. Thus, non-canonical ubiquitylation might have more far-reaching biological impacts than previously anticipated.

## KEY RESOURCES TABLE

| REAGENT or RESOURCE | SOURCE | IDENTIFIER |
| --- | --- | --- |
| Bacterial Strains |  |  |
| BL21(DE3) cells | New England Biolabs | Cat# C2527H |
| Chemicals, Peptides, and Recombinant Proteins |  |  |
| Glutathione Sepharose 4B | Expedeon | Cat# AGSCUST |
| Ubiquitin | In house production |  |
| Nucleophiles |  |  |
| Deposited Data |  |  |
| Software and Algorithms |  |  |
| Prism | GraphPad | https://www.graphpad.com/scientific-software/prism/ |
| COOT | Emsley et al., 2010 | http://www2.mrc-lmb.cam.ac.uk/personal/pemsley/coot/ |
| PyMOL |  | https://pymol.org/2/ |
| Adobe Illustrator |  | https://www.adobe.com/uk/products/illustrator.html |

cDNA constructs and proteins
| Protein | Expressed protein | Tag<br>Cleaved | Vector<br>type | Plasmid | DU number |
| --- | --- | --- | --- | --- | --- |
| UBE2Q1<sup>FL</sup> | GST- | Yes | Bacterial | pGEX6P1 | 4213 |
| UBE2Q1<sup>cat</sup> | GST-UBE2Q1 220-422 (end) (WT) | Yes | Bacterial | pGEX6P1 | 23956 |
| UBE2Q1<sup>cat</sup> | GST-3C-UBE2Q1 D220-G422 C351S | Yes | Bacterial | pGEX6P1 | 61423 |
| UBE2Q1<sup>cat</sup> | GST-3C-UBE2Q1 Y343A D220-G422 | Yes | Bacterial | pGEX6P1 | 73951 |
| UBE2Q1<sup>cat</sup> | GST-3C-UBE2Q1 L345A D220-G422 | Yes | Bacterial | pGEX6P1 | 73272 |
| UBE2Q1<sup>cat</sup> | GST-3C-UBE2Q1 L405A D220-G422 | Yes | Bacterial | pGEX6P1 | 73273 |
| UBE2Q1<sup>cat</sup> | GST-3C-UBE2Q1 I408A D220-G422 | Yes | Bacterial | pGEX6P1 | 73274 |
| UBE2Q1<sup>cat</sup> | GST-3C-UBE2Q1 H409A D220-G422 | Yes | Bacterial | pGEX6P1 | 73264 |
| UBE2Q1<sup>cat</sup> | GST-3C-UBE2Q1 H409N D220-G422 | Yes | Bacterial | pGEX6P1 | 73955 |
| UBE2Q1<sup>cat</sup> | GST-3C-UBE2Q1 N412A D220-G422 | Yes | Bacterial | pGEX6P1 | 73263 |
| UBE2Q1<sup>cat</sup> | GST-3C-UBE2Q1 W414A D220-G422 | Yes | Bacterial | pGEX6P1 | 73277 |
| UBE2Q1<sup>cat</sup> | GST-3C-UBE2Q1 W414F D220-G422 | Yes | Bacterial | pGEX6P1 | 73954 |
| UBE2Q1<sup>cat</sup> | GST-3C-UBE2Q1 W414Q D220-G422 | Yes | Bacterial | pGEX6P1 | 73975 |
| UBE2Q1<sup>cat</sup> | GST-3C-UBE2Q1 Y415A D220-G422 | Yes | Bacterial | pGEX6P1 | 73276 |
| UBE2Q1<sup>cat</sup> | GST-3C-UBE2Q1 P417G D220-G422 | Yes | Bacterial | pGEX6P1 | 73945 |
| UBE2Q1<sup>cat</sup> | GST-3C-UBE2Q1 P417A D220-G422 | Yes | Bacterial | pGEX6P1 | 73946 |
| UBE2Q1<sup>cat</sup> | GST-3C-UBE2Q1 P418A D220-G422 | Yes | Bacterial | pGEX6P1 | 73270 |
| UBE2Q1<sup>cat</sup> | GST-3C-UBE2Q1 P418G D220-G422 | Yes | Bacterial | pGEX6P1 | 73271 |
| UBE2Q1<sup>cat</sup> | GST-3C-UBE2Q1 D421A D220-G422 | Yes | Bacterial | pGEX6P1 | 73275 |
| UBE2Q1<sup>cat</sup> | GST-3C-UBE2Q1 D421L D220-G422 | Yes | Bacterial | pGEX6P1 | 73952 |
| UBE2J2 | GST 3C UBE2J2 T11-T183 | Yes | Bacterial | pGEX6P1 | 61848 |
| UBE2J2 | GST 3C UBE2J2 T11-T183 D99A | Yes | Bacterial | pGEX6P1 | 72870 |
| UBE2J2 | GST 3C UBE2J2 T11-T183 F100A | Yes | Bacterial | pGEX6P1 | 72894 |
| UBE2J2 | GST 3C UBE2J2 T11-T183 H101A | Yes | Bacterial | pGEX6P1 | 72871 |
| UBE1 | His-TEV-UBE1 | Yes | Insect | pFastBac | 32888 |
| Ubiquitin 1-76 | Ubiquitin (expressed tagless) | Tagless | Bacterial | pET24 | 20027 |

## RESOURCE AVAILABILITY

### Lead contact

Further information and requests for resources and reagents should be directed to and will be fulfilled by the Lead Contact: Virginia De Cesare

### Materials Availability

Plasmids used in this study have been deposited with and will be distributed by MRC PPU reagents and services (https://mrcppureagents.dundee.ac.uk/)

## Acknowledgments

We thank Prof. Ron Hay, Prof. Satpal Virdee, Prof. Helen Walden and Prof. Dario Alessi for useful discussions. We thank the Medical Research Council (MRC) Antibody Development team for their support in raising anti-UBE2Q1 antibody. We thank the MRC PPU Reagents and Services Antibody Development team (https://mrcppureagents.dundee.ac.uk/) for their support in raising anti-UBE2Q1 antibody. This work was funded by UKRI (Grant Reference MR/V025759/1). We also acknowledge pharmaceutical companies supporting the Division of Signal Transduction Therapy (Boehringer-Ingelheim, GlaxoSmithKline, and Merck KGaA).

## Supplementary Figures

**Figure S1.**
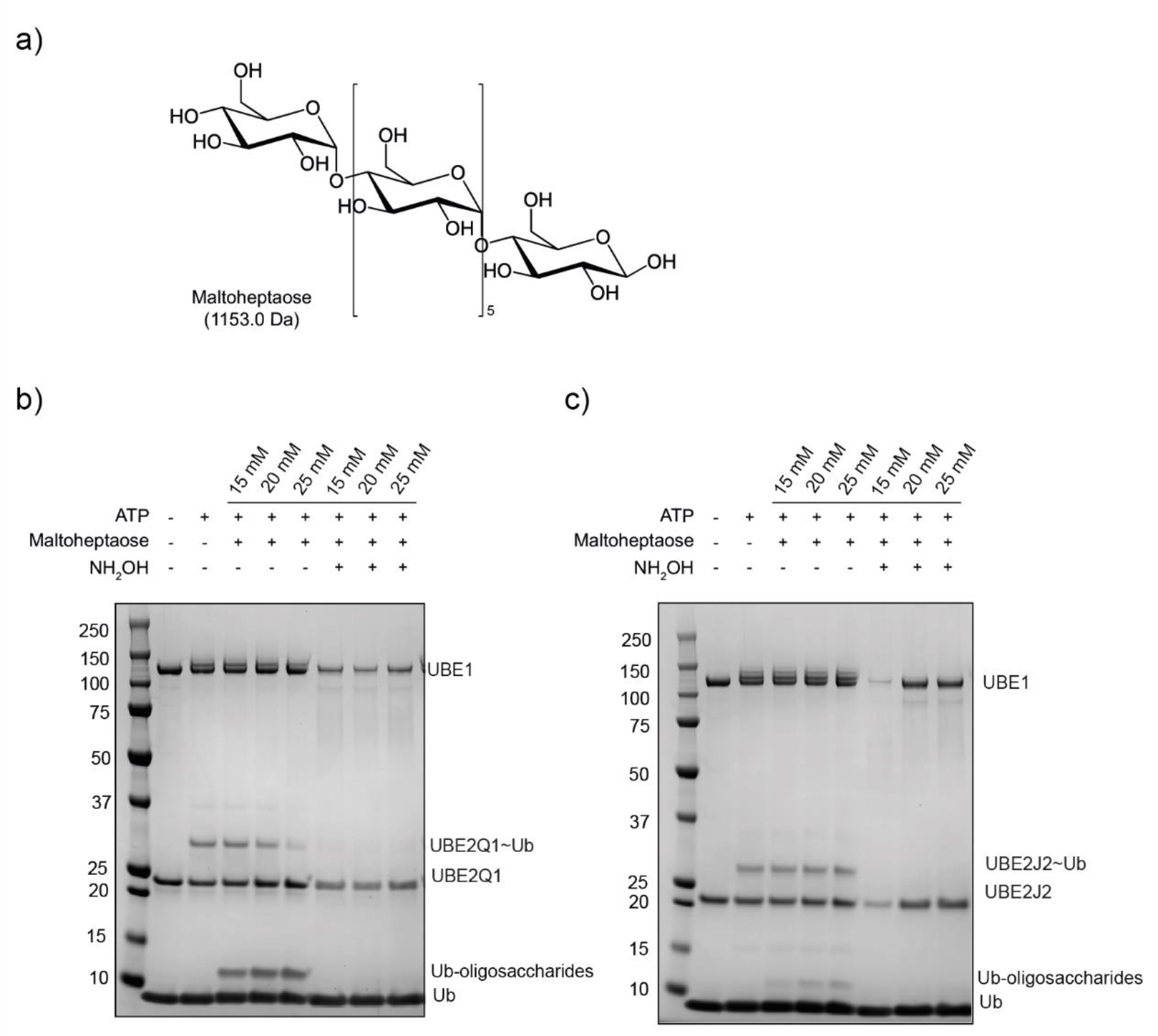
UBE2Q1 and UBE2J2 ubiquitylate malthoheptaose. Structure of maltoheptaose (a). UBE2Q1 (b) or UBE2J2 (c) directly ubiquitylated maltohepatose.

**Figure S2.**
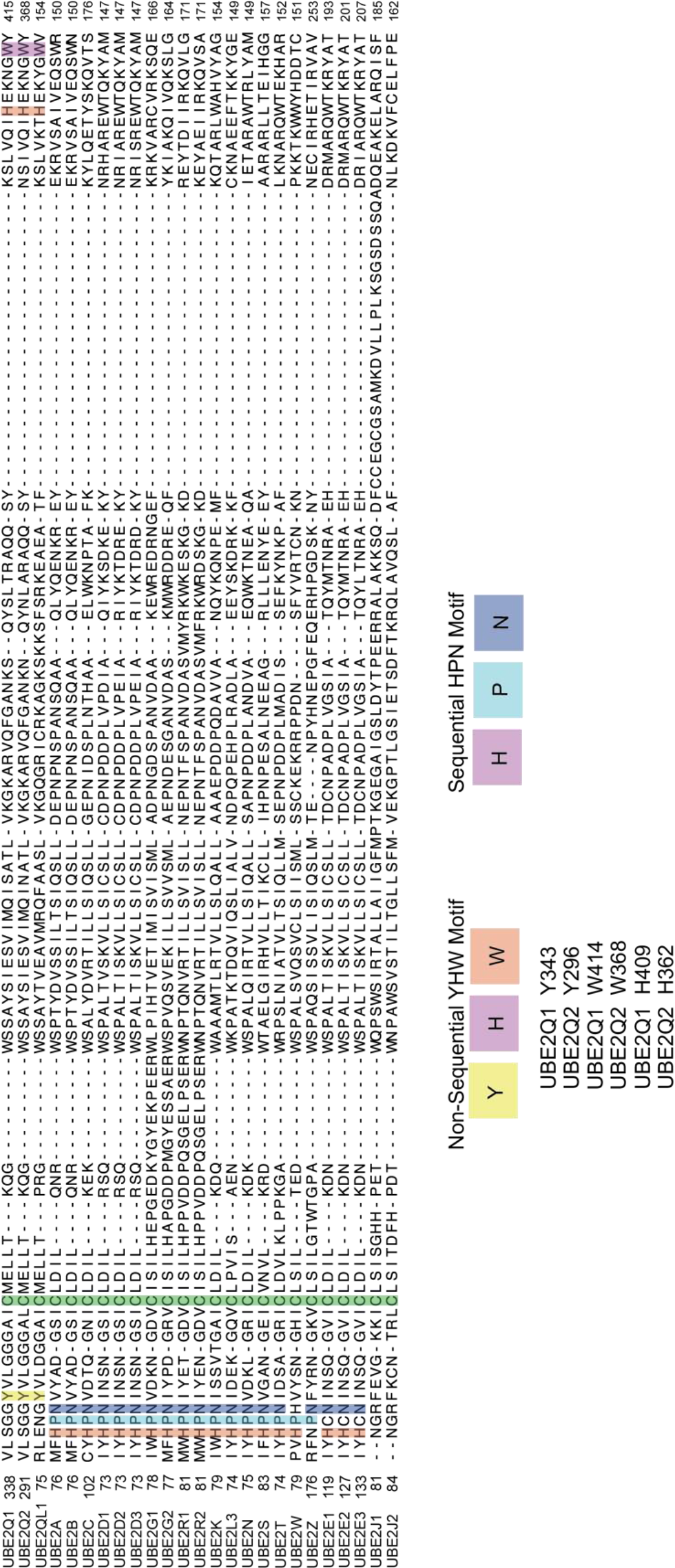
Alignment of the UBC domain of 28 ubiquitin E2 conjugating enzymes reveals absence of the canonical HPN motif in the UBE2Q and UBE2J families. Highlighted UBE2Q1, UBE2Q2 and UBE2QL1 non-sequential activity determinants (YHW motif) and the canonical HPN triad. Catalytic cysteine marked in green.

**Figure S3.**
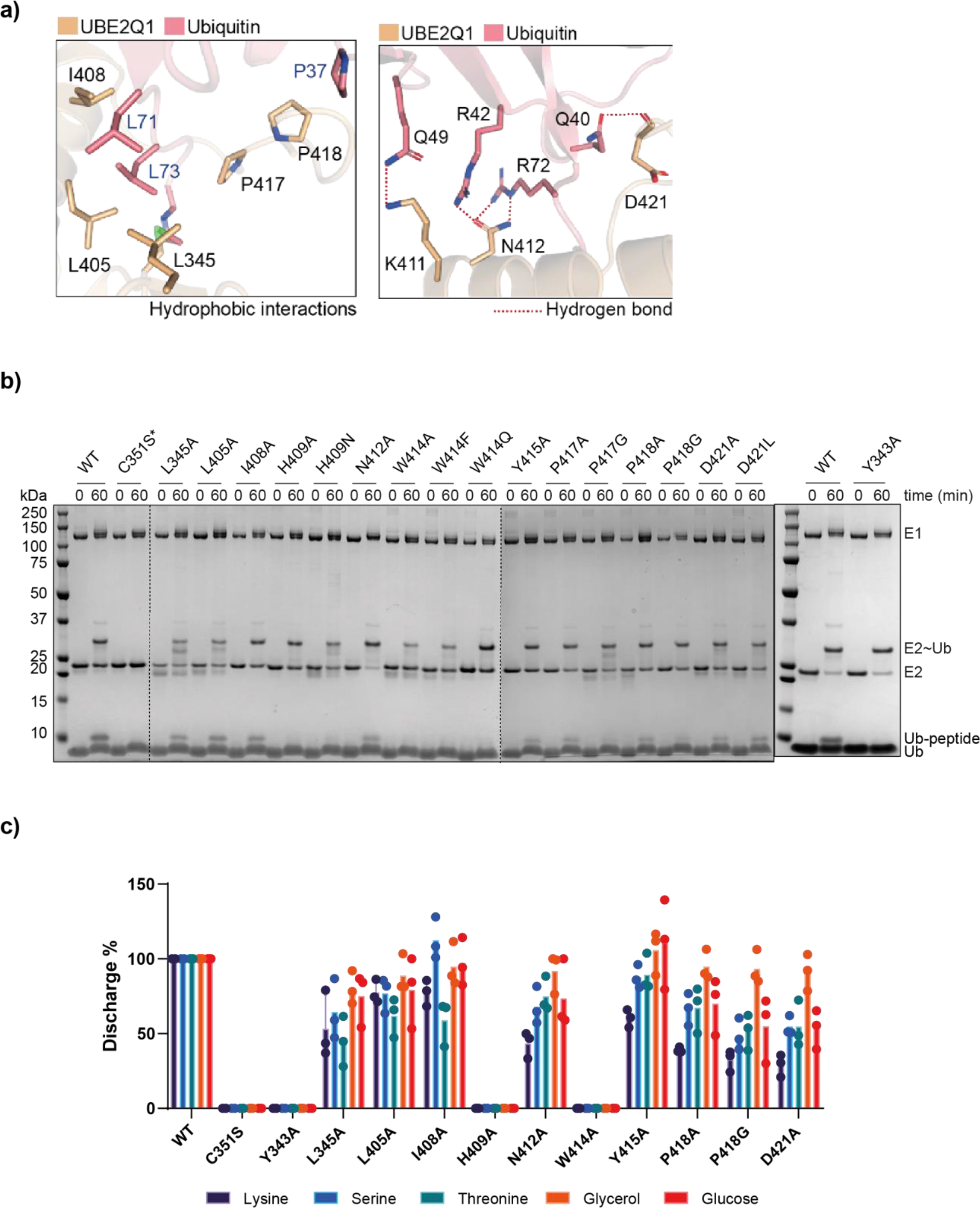
Structural model of the UBE2Q1∼Ub interaction. Hydrophobic and Hydrogen bond- based interactions reported. Indicated UBE2Q1 mutants were tested for their ability to ubiquitylate B4GALT1 peptide 1 (b) and to discharge on the indicated nucleophiles by MALDI-TOF MS (c).

**Figure S4.**
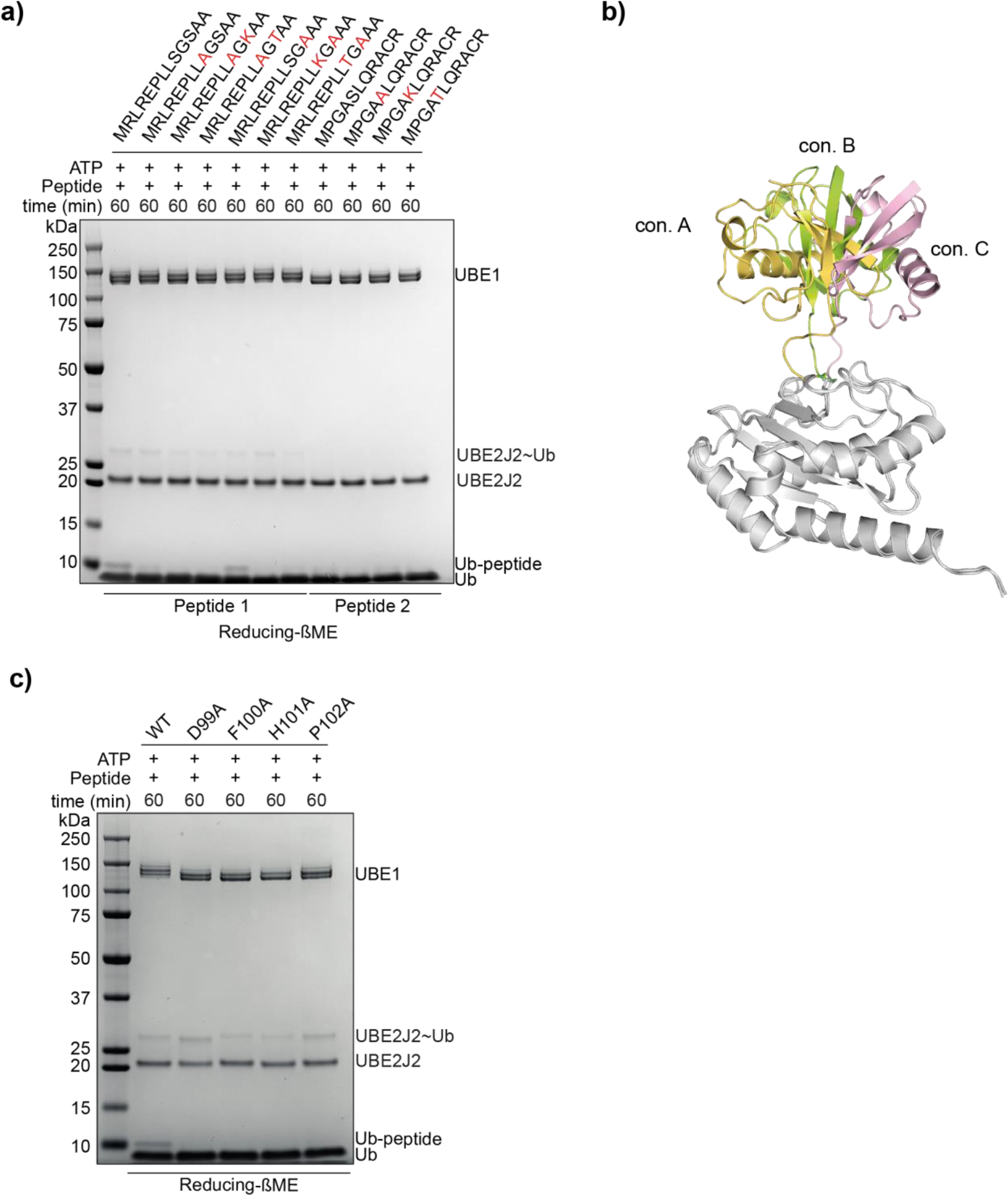
UBE2J2 ubiquitylates B4GALT1 CD on B4GALT1 peptide 1 and peptide 2 (a): reducing conditions completely abolish cysteine based ubiquitylation on peptide 2. UBE2J2 mutations that affect the discharge on B4GALT1 peptide 1, reducing conditions (b).

**Figure S5.**
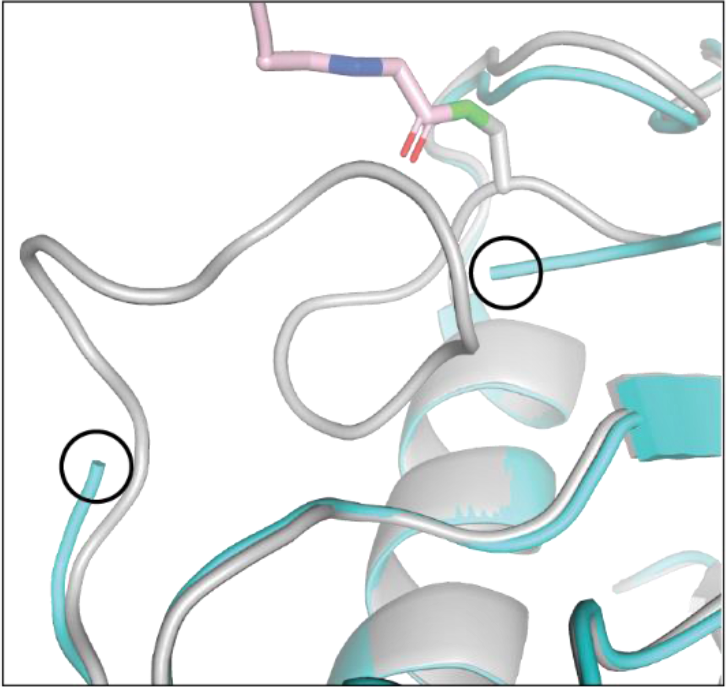
UBE2J2 apo crystal structure (reported in light blue) and predicted conformation of the UBE2J2 highly mobile stretch of residues interacting with ubiquitin (reported in grey)

**Figure S6.**
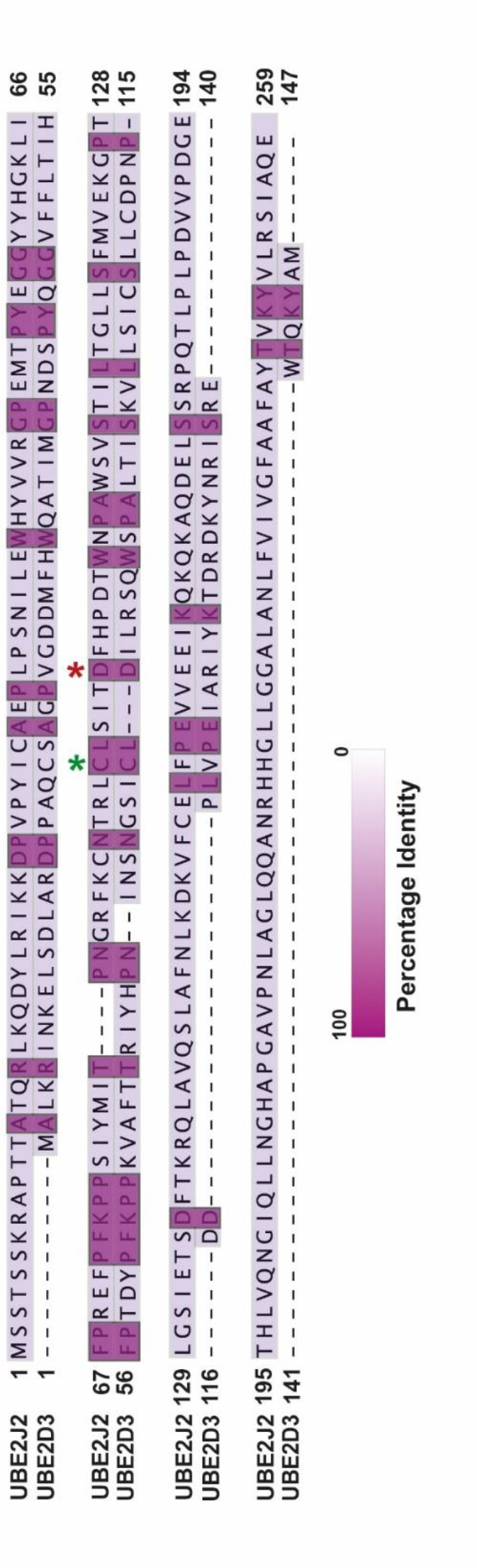
UBE2D3 and UBE2J2 alignment, catalytic cysteine indicated with green asterisk. UBE2J2 D99 residue aligns with the D87 residue in the canonical UBE2D3 E2 conjugating enzyme (red asterisk).

## Notes

### Competing Interest Statement

The authors have declared no competing interest.

